# Porcupine: a visual pipeline tool for neuroimaging analysis

**DOI:** 10.1101/187344

**Authors:** Tim van Mourik, Lukas Snoek, Tomas Knapen, David Norris

## Abstract

The field of neuroimaging is rapidly adopting a more reproducible approach to data acquisition and analysis. Data structures and formats are being standardised and data analyses are getting more automated. However, as data analysis becomes more complicated, researchers often have to write longer analysis scripts, spanning different tools across multiple programming languages. This makes it more difficult to share or recreate code, reducing the reproducibility of the analysis. We present a tool, Porcupine, that constructs one’s analysis visually and automatically produces analysis code. The graphical representation improves understanding of the performed analysis, while retaining the flexibility of modifying the produced code manually to custom needs. Not only does Porcupine produce the analysis code, it also creates a shareable environment for running the code, in the form of a Docker image. Together, this forms a reproducible way of constructing, visualising and sharing one’s analysis. Currently, Porcupine links to Nipype functionalities, which in turn accesses most standard neuroimaging analysis tools. With Porcupine, we bridge the gap between a conceptual and an implementational level of analysis and thus create reproducible and shareable science. We give the researcher a better oversight of their pipeline, both while developing and communicating their work. This will reduce the threshold at which less expert users can generate reusable pipelines. We provide a wide range of examples and documentation, as well as installer files for all platforms on our website: https://timvanmourik.github.io/Porcupine. Porcupine is free, open source, and released under the GNU General Public License v3.0.

**Author Summary:** The neuroimaging community is fervently debating that its reproducibility and transparency should be improved, but it is a challenging problem as to how to accomplish this. We here propose a tool, Porcupine, to aid in this process by more easily creating shareable workflows for analysing neuroimaging data. The conceptual understanding of a pipeline is improved by means of the graphical interface, and it automatically produces the code to perform the analysis and to create a sharing environment. This retains full flexibility to modify the script afterwards but in principle produces readily executable code for an end-to-end analysis. Porcupine currently links to all Nipype functionality, but is designed to be extendable to other workflow packages in neuroimaging and beyond. Porcupine is free and is released under the GNU General Public License.

## Introduction

The field of neuroimaging is rapidly adopting a more reproducible approach to data acquisition and analysis. Especially in recent years, a strong movement for conducting better documented and more reproducible science can be observed. Advances have been made in terms of openly sharing data (e.g. OpenFmri, [1]), standardizing data formats (BIDS format [2]), and facilitating more automated pipelines [3–5]. These initiatives facilitate increasing global scientific communication and collaboration, that is paramount in the age of big data.

As a result of the increasing complexity of analyses, however, and the wide variety of different tools, researchers often have to write custom scripts for combining different software packages, often in different programming languages. As an extra obstacle, many tools have external dependencies, intricate installation procedures, or different file formats for the same type of data. Furthermore, the sharing initiatives usually have a stronger focus on sharing *data* (Human Connectome Project [6], NeuroVault [7]) instead of *code*, such that analysis scripts still have to be recreated based on the method section of a paper. All these factors negatively affect the reproducibility, documentation, and in the worst case correctness of the analysis [8].

A considerable mastery of coding is required for analysing fMRI data. The conceptual side of understanding all preprocessing steps is not trivial, but converting this into a working pipeline can be an arduous journey. The necessary programming skills are not usually the prime focus of a brain researcher’s skills or interests, but they are a necessity for completing one’s analysis. Consequently, scripting a pipeline that covers all high-level and low-level aspects is daunting and error prone. As a result, there is a considerable risk one will revert to ‘hacking’ an analysis pipeline together, sacrificing a reproducible approach. So as a researcher, how do you start an analysis? It is easiest to start with visualising the steps of your analysis pipeline.

In an increasingly complicated analysis environment there is a strong need for tools that give a better oversight of these complex analyses, while retaining the flexibility of combining different tools. A notable effort to integrate different tools is Nipype [4], that has a Python interface to existing tools from all major MRI analysis packages. However, this still requires non-trivial Python scripting. Furthermore, Nipype is only able to visualise a workflow after it has been manually scripted [9].

Here we detail our solution to these problems, an open-source software program we call Porcupine: ‘PORcupine Creates Ur PipelINE - the worst recursive acronym with bad capitalisation and annoying use of slang.’ Porcupine allows the creation of neuroimaging pipelines by means of a graphical user interface (GUI). After graphical pipeline definition, Porcupine in turn creates the code that programmatically defines the pipeline. Additionally and without any additional overhead, we supply a Dockerfile (https://www.docker.com) that automatically builds the run environment for the pipeline. This not only facilitates sharing the pipeline, but also ensures its reproducibility [10]. We provide an extensive list of examples and documentation on our website, as well as the possibility to upload one’s custom pipeline to create a community driven library of analyses.

By implementing an intermediate visual step in the generation of preprocessing workflows, Porcupine allows the user to focus on the logical flow of the preprocessing pipeline in a graphical representation without the need for coding at this conceptual stage of development. Because the GUI produces working analysis code the user can then immediately inspect, save, and run the generated code. Thus, Porcupine provides a stepping stone that eases the transition from concept to implementation. Because the entire pipeline and its parameters are defined *in abstracto* before it is run, systems such as Nipype allow for elaborate checks and optimisations of the pipeline’s execution. Furthermore, such systems can straightforwardly incorporate full logging of all analysis steps, creating a paper trail of the pipeline’s execution. This combination of a reproducible environment in which a predefined pipeline is run by means of a system that provides precise bookkeeping paves the way to new standard that will ensure steady and reproducible progress in the field of cognitive neuroimaging [11].

In our practical experience, the use of Porcupine allows one to very quickly prototype preprocessing pipelines. Novice users can create a pipeline *de novo* and quickly focus on the code for this pipeline, greatly speeding up the learning process and thereby facilitating the use of reproducible pipelines. We envisage Porcupine to play a role in both the education of novice neuroimaging students and the rapid prototyping of pipelines by expert users. Here, we first outline several Porcupine use-case scenarios of increasing complexity, after which we detail the architecture of Porcupine.

## Results

### What is Porcupine?

Porcupine is a graphical workflow editor that automatically produces analysis code from a graphically composed pipeline. By dropping ‘nodes’ (representing analysis steps) into the workflow editor and by connecting their data inputs and outputs, a pipeline is constructed. Analysis code is then automatically generated from the graphical representation of the pipeline. The code can readily be saved to a script (e.g. a Python, MATLAB, or Docker file) in order to perform the desired analysis. Additionally, the pipeline can be shared or inspected in visual form (PDF/SVG), or saved to a Porcupine specific (.pork) file to continue working on the pipeline at another time.

Apart from the visual representation of the pipeline, we provide more functionality to orderly structure one’s analysis, as outlined in Fig. 1. All functions (the nodes in the graph) that are included in the pipeline are also listed in a separate panel, listing their input parameters, output data, as well as a link to the online documentation of the function. We also provide the option to iterate over any input variable in order to facilitate parallelisation over subjects, sessions, or other variables. All parameters may also be edited in a separate parameter panel of the user interface. This functions as a central storage for important parameters, for example the ones that should be reported in a methods section. Porcupine combines the graphical overview and the parameters to automatically create the analysis code shown in the code window.

We here focus on code generation that strictly adheres to the Nipype API [4], a Python-based MRI analysis and pipelining package. Nipype is used for its strong focus on uniformity in accessing functions, its link to most major MRI analysis tools, and its emphasis on reproducible science. Porcupine’s architecture, however, is in principle agnostic with respect to the specific implementation of the underlying pipelining software. Any package with a consistent interface in the field of e.g. neuroimaging, bioengineering, or astronomy could benefit from using Porcupine’s architecture.

We first show that we can easily generate a standard fMRI analysis pipeline. After visually dragging and dropping modules, code is automatically created that is usually scripted manually instead. We then show how we facilitate loading data from an online repository, generate a readily executable fMRI pipeline, but also generate a shareable and reproducible analysis environment (using Docker), all with minimal additional effort. This allows for easily scalable analyses that could be performed locally, but also on computational clusters or with cloud computing, without manual installation of different software packages.

**Fig. 1.**
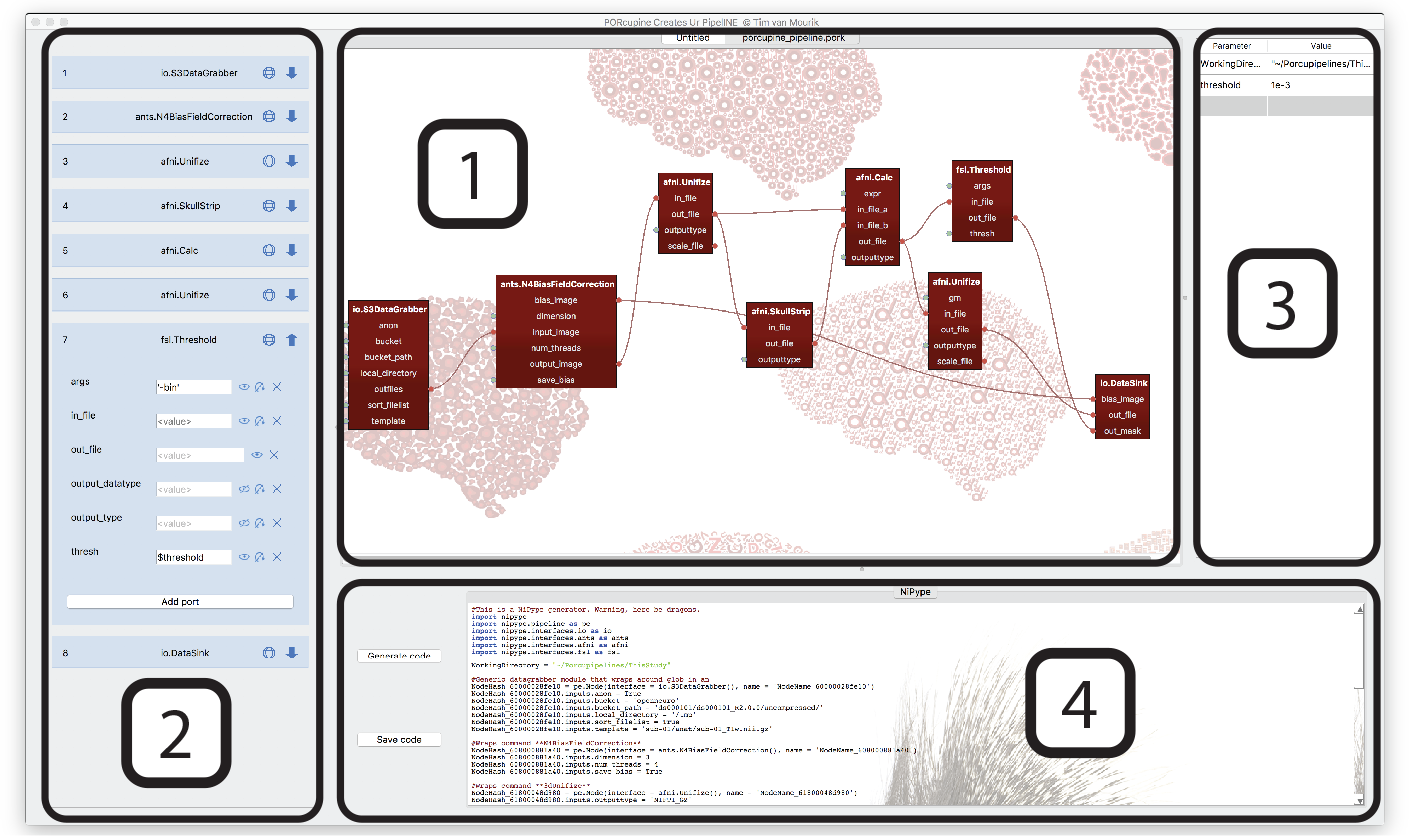
A screenshot of a Porcupine workflow. The editor is divided into four panels, each of them targeted at facilitating a more understandable and reproducible analysis. The *workflow editor* (1) provides a visual overview of one’s analysis. The functions are all listed in the *node editor* (2), where the parameters for all functions can be orderly stored. This may include links to important parameters that are listed in the *parameter editor* (3), such that an overview of the main analysis settings can be easily viewed and modified. Readily executable analysis code is generated in the *code window*(4)

## Usage example

We here show a simple example that constructs a pipeline for a single operation. In three steps, data is loaded, (minimally) processed, and the output is written to disk, as shown in Fig. 2. We here show an example that links to an OpenNeuro fMRI data set,but we could load any online data set that is set up according to the BIDS format [2]. OpenNeuro’s data sets are stored as Amazon repositories (‘S3 buckets’) and can be loaded by dragging the appropriate module into the workflow editor and typing the name of the bucket into the node editor. Its output can subsequently be connected to a Nipype function node, for example FSL’s Brain Extraction Tool. All parameters of the function are listed and can be set in two different ways: either by dragging a link from a previous node’s output port to an input port in the next node, or by typing in the parameter in the node editor. Subsequently, output can be written to disk by connecting the desired output to a Nipype DataSink node that collects and stores the data. By pressing the ‘Generate code’ button, the code for this pipeline is automatically generated and can immediately be saved and executed in a Python shell.

## Pipeline sharing

From a simple example that reads and writes the data, a more complicated pipeline is readily set up. More functionality, i.e. nodes, can be dragged in and connected to quickly build a custom pipeline. As it is commonplace to repeat a single analysis or function for several subjects, sessions, or other variables, every field can be flagged as an ‘iterator’ field. This facilitates looping over variables. Once the pipeline is set up and the code is generated, Nipype offers functionality to construct a visual pipeline graph from custom python code. In Porcupine’s proposed use-case, this end point of a standard Nipype pipeline represents the starting point, as shown in Fig. 3. This allows the user to focus on the desired pipeline graph first, and then progress to the manual editing of the generated code.

**Fig. 2.**
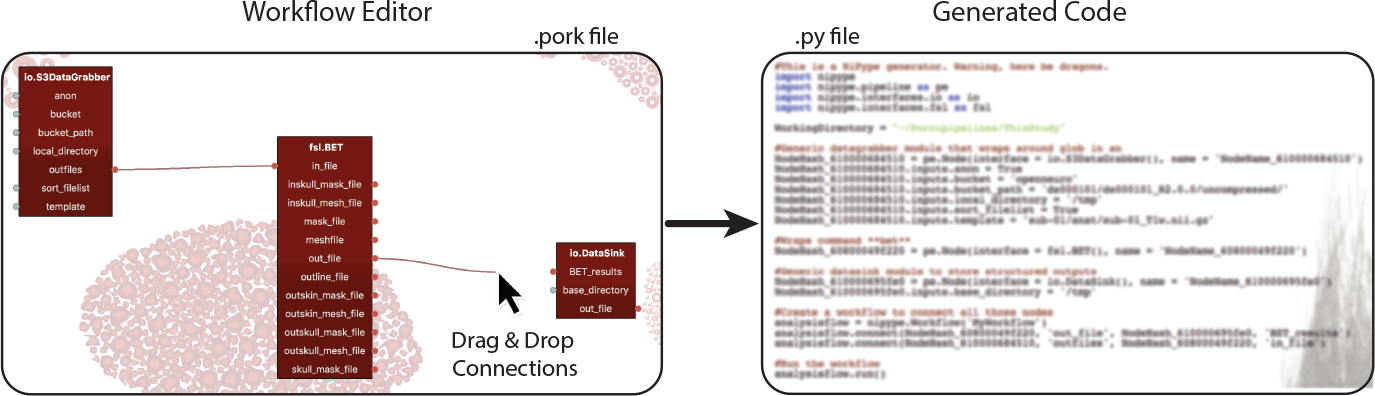
An example of simple workflow. In three steps, this pipeline loads data, processes it, and writes it to disk. This is achieved by connecting the input and output fields from subsequent nodes in the pipeline. The constructed workflow is then transformed in readily executable (Nipype) analysis code.

**Fig. 3.**
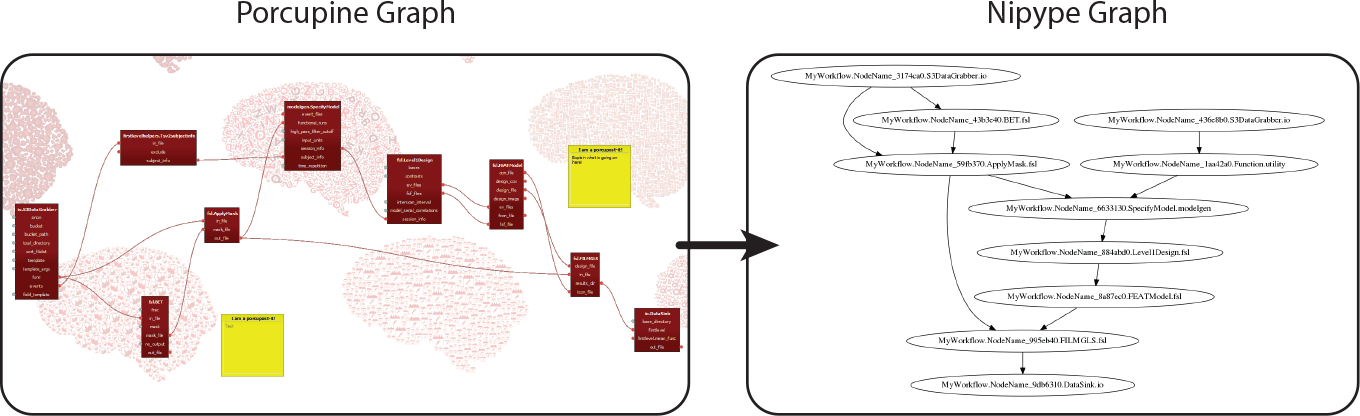
An example of a more complicated and realistic fMRI preprocessing pipeline. Once the code is generated, this can in turn be transformed into a Nipype graph visualisation. Whereas this is usually the end point for a pipeline in Nipype, we here propose to use a visualisation as a starting point of one’s analysis.

Not only does Porcupine provide a way of setting up a preprocessing or analysis pipeline, we also provide a means for executing these pipelines in a reproducible environment. In addition to the Python analysis file that is generated, we create a scaffold for a Docker file. Docker <u>(https://www.docker.com)</u> is an open platform to easily build, run and share applications. The generated Docker file describes a minimal operating system that is required to run the analysis, based on the dependencies of the modules used in the workflow editor. With this Docker file, an image of the full analysis can be built, shared and executed. This provides a simple platform to reproduce results of one’s analysis, on the same data set, or on another with only a single change in the data source module. Alternatively, one can use it as a template environment for a new follow-up analysis. As with all generated code, the Docker code is fully customisable to a researcher’s need, but our suggested scaffold requires only a single manual edit to be built as a Docker image (see S1 Docker files). The Docker command will execute the pipeline: load the data from an online repository, process the data, and store only the output data to a local directory. The Docker image includes both the pipeline code and the run environment, and can be shared alongside a paper via DockerHub. The above examples (and many more) as well as extensive documentation and tutorials can be found here.

## Limitations

Some features in Nipype have not been implemented. Notably, the JoinNode functionality is not yet accessible from the Porcupine user interface, in which the results from an upstream iterator are aggregated to a single output. Furthermore, custom extensions of Nipype functions are not automatically supported, but we do provide a script to add one’s own custom module to Porcupine that would make this functionality accessible. A GUI for this is still an intended point of improvement. In general, feature requests are maintained as *projects* in the GitHub repository. We encourage people to contribute new ideas or implementations for functionality in terms of modules, new toolboxes, and, most importantly, custom pipelines that can be added to the repository. Details for contributing can be found on the website.

While Porcupine in principle supports all workflow operations, a specific pipeline may well require modules that are not provided within Nipype. It is advised that the user either packages custom code for this into a new module, or manually adds it to the produced code. Porcupine itself functions only as the front-end to the Nipype back-end, but does not perform type-matching on the input and output of a connection, nor does it perform syntax checking of the manually edited parameters.

## Methods

Porcupine’s graphical user interface was written first with a general visual programming application in mind. The initial interface to Nipype was developed at a three-day coding sprint at BrainHack 2017, Amsterdam. This kickstarted Porcupine in its current form. The source code, as well as the installer files for Windows, Mac, and Linux, are publicly available as a GitHub repository. Porcupine is free, open source, and released under the GNU General Public License v3.0.

Visual programming is a generic way of programming to create a data flow or to perform an ordered task with a modular structure [12]. Customarily, it allows the user to construct a Directed Acyclic Graph (DAG) [13] of conceptualised operations that are subsequently interpreted or compiled as an application [14]. This format is particularly useful for workflows that fit modular structures, such as most neuroimaging data analyses [15].

## Architecture

Not only do we intend researchers to make their analyses (re-)usable and robust, our software also adheres to all 20 simple rules that were laid out to this end [16,17]. The updates as well as the releases of the source code are realised by means of a GitHub repository. Installer files are provided for all platforms and do not require administrator privilege. Users are aided in getting started quickly by extensive documentation and an example gallery.

Easy cross-platform installation or compilation was achieved by programming Porcupine as a stand-alone application in Qt Creator (https://www.qt.io) for C++. Internal file formats were standardised to JSON dictionaries, a format native to Python, Qt, and web applications. This provides a simple means to add new modules to Porcupine, without the need to write additional code. Every dictionary specifies a software package (e.g.‘Nipype’,‘Docker’, etc.) that is interpreted by Porcupine and creates code that is native to the package. A package-specific interpreter needs to be written just once, after which new modules that are included in the dictionary will be automatically available in Porcupine.

Each JSON dictionary describes a list of functions (internally referred to as ‘nodes’). Each function has a name and (optionally) a category, a web url to its documentation, and a block of code. A code block specifies the software package for which the node is meant, the associated piece of code for that function and optionally an additional comment. Furthermore, a node contains any number of data/parameter ports, each of which can be input, output, or both. Optionally, additional flags can be set for ports to be visible in the editor, whether its value is editable, or whether the variable needs to be iterated over. Thus, JSON files for custom nodes can easily be created and added as a dictionary to the graphical interface. We also provide a Python script that converts a custom Python function(s) to a Nipype node dictionary.

## Extending Porcupine with new toolboxes

Currently, Porcupine features Nipype and Docker support, but this could easily be extended to other software packages. This requires no major changes to the Porcupine source code, merely the inclusion of a single C++ class that describes the relationship between the nodes, links, and the output code. Specifically, the ‘CodeGenerator’ class must be inherited and has access to the full workflow: the list of nodes, their parameters, and their connections. As long as all functions within an analysis toolbox can be accessed with a consistent interface, they can be represented as modules within Porcupine. Apart from Nipype, support for a laminar specific fMRI analysis toolbox in MATLAB is provided. The developers of the Fastr framework programmed initial support for their code base [18]. Unfortunately, only few neuroimaging packages abide by this uniformity of their functions and hence many cannot be included into Porcupine.

## >Relation to existing pipeline managers

Porcupine aims to provide an extendable, transparent and flexible platform to build preprocessing and analysis pipelines. Other software packages have made similar attempts at providing visual aids to build or run pipelines. Within neuroimaging, the most notable ones are the JIST pipeline [19], extended with CBS Tools [20] and the LONI pipeline [15]. Porcupine distinguishes itself from these by not creating a run environment, but instead creating the analysis code for the researcher. This retains the possibility of immediately running the code through a Python interpreter, but also creates more flexibility, as researchers can modify and adjust the script according to their needs. Additionally, as the functions link to existing Nipype interfaces, error reporting is more transparent such that problems can more easily be resolved. Lastly, our open-source framework is set up to be extendable with new modules within existing frameworks, as well as with completely new frameworks. This provides a future-proof set-up for current and future analysis tools in neuroimaging and perhaps other disciplines.

## Discussion

We have presented a new tool to visually construct an analysis pipeline. Subsequently, Porcupine automatically generates the analysis code, and provides a way of running and sharing such analyses. We see this as an important tool and a stepping stone on the path to doing more reproducible and open science. Additionally, this gives researchers a better oversight of their analysis pipeline, allowing for greater ease of developing, understanding, and communicating complex analyses.

Porcupine provides two independent functionalities that dovetail to allow users to more easily take part in reproducible neuroimaging research. They are (1) a graphical user interface for the visual design of analysis pipelines and (2) a framework for the automated creation of docker images to execute and share the designed analysis.

We anticipate that the ability to design processing pipelines visually instead of programmatically cuts the novice user’s learning phase by a considerable amount of time by facilitating understanding and development. The ease of use of a Graphical User Interface (GUI) implementation extends and complements Nipype’s flexibility. Thus, it invites researchers to mix and match different tools, and adhere less stringently to the exclusive use of the tools of any given toolbox ecosystem. This flexibility enhances the possible sophistication of processing pipelines, and could for instance be helpful in cross-modal research or multi-site research. Additionally, it may nudge method developers to write new tools in a way that easily integrates with the Nipype and Porcupine structure.

The emphasis that Porcupine puts on visual development of analyses makes it easier to communicate a methods section visually rather than in writing. We foresee that researchers may prefer explicity sharing the created .pork files and the Nipype pipelines that are created from them, instead of solely relying on written descriptions of their methods. Yet another use case for Porcupine is the easy definition of proposed processing workflows for preregistered studies.

Importantly, Porcupine attempts to reduce the steepness of the learning curve that is inherent to the use of complex analysis, by providing a more structured and systematic approach to pipeline creation. It separates the skill of building a conceptual analysis pipeline from the skill of coding this in the appropriate programming language. This places Porcupine in a position to aid in the education of novice neuroimaging researchers, as it allows them to focus on the logic of their processing instead of creation of the code for the processing - greatly improving and accelerating their understanding of the different steps involved in the preprocessing of neuroimaging data. At the same time, it allows more experienced researchers to spend more time on theconceptual side than on implementational side.

Having allowed for the visual design of a pipeline for the preprocessing or analysis of a neuroimaging dataset, the reproducible execution of this pipeline is another step that Porcupine facilitates. By flexibly creating a Docker image tailored to the different preprocessing steps defined visually in the GUI, Porcupine allows the user to share not only the definition of the pipeline but also its execution environment. This step removes the overhead of having to manually install the desired operating system with the matching distribution of MRI analysis software. This final step greatly facilitates the reproducibility of reported results, and is part of a general evolution of the field towards easily shareable and repeatable analyses.

The generated Docker image can be made High Performance Computing aware with singularity by means of dedicated tools such as docker2singularity. Alternatively, with only trivial additions to the Dockerfile, it can be transformed into a BIDS app [21]. A detailed explanation for doing this can be found on our website. An automatic and direct way of creating has not yet been implemented. Additionally, integrating support for standardised workflow file formats, such as the Common Workflow Language [22] could further add to Porcupine’s aim of reproducibility. Another point of improvement is a functionality to embed pipelines within pipelines. Currently, a complicated pipelines does full justice to the term ‘spaghetti code’, and the number of nodes and links may easily compromise the visual aid in understanding; the very purpose for which Porcupine was created. This may easily be solved by compartmentalising pipelines into logical units by providing an embedded structure.

We intend Porcupine to be a strong aid for doing better, more reproducible and shareable science. By bridging the gap between a conceptual and implementational level of the analysis, we give scientists a better oversight of their pipeline and aid them in developing and communicating their work. We provide extensive and intuitive documentation and a wide range of examples to give users a frictionless start to use Porcupine. We look forward to adding more functionality and support for more toolboxes in the near future.

## Supporting information

**S1 Docker files** Porcupine provides a Docker image that creates the necessary run time environment for a pipeline that is constructed in the workflow editor. As with all generated code, the Docker code is fully customisable to a researcher’s need, but our suggested scaffold requires only a single manual edit to be built as a Docker image. A Docker script can only refer to online or on-disk resources, so the pipeline file needs to be saved manually and added to the Docker file:

~~~
ADD / path /to/ pipeline / script .py / somewhere / porcupipeline .py 298
CMD [" python ", "/ somewhere / porcupipeline .py"]
~~~

Once this line is added, the docker image can be built:

~~~
$ docker build -t mydockerimage -f Dockerfile
~~~

The output from the executed pipeline can be written to a local directory on a researcher’s computer (‘/my/local/directory’) by mounting it to the docker output directory (‘/data’) with the ‘-v’ option and running the image as if it were a standalone application.

~~~
$ docker run -v /my/local/directory :/ data mydockerimage
~~~

This Docker command will execute the pipeline: load the data from an online repository, process the data, and store only the output data to a local directory. A fully worked out example with detailed explanation can be found here.

## Acknowledgements

We would like to thank the organisation of BrainHack Global and BrainHack Amsterdam, specifically Pierre-Louis Bazin, for organising the platform that kickstarted Porcupine in its current form.

## Author contributions

TvM wrote the Porcupine C++ software. TvM, LS, and TK designed the interface to Nipype. LS and TvM built the website. LS wrote the examples and the majority of documentation. TvM, TK, LS, and DGN wrote the paper.

